# Lower limb chronic edema management program: Perspectives of disengaged patients on challenges, enablers and barriers to program attendance and adherence

**DOI:** 10.1101/693135

**Authors:** Linda A. M. Khong, Amma Buckley, Wendy Johnson, Vinicius Cavalheri

**Affiliations:** Physiotherapy Department, Sir Charles Gairdner Hospital, Nedlands, Western Australia, Australia; Institute for Health Research, University of Notre Dame Australia, Fremantle, Western Australia, Australia; Media, Cultural Studies and Social Inquiry, Curtin University, Perth, Western Australia, Australia; School of Physiotherapy and Exercise Science, Faculty of Health Sciences, Curtin University, Perth, Western Australia, Australia

## Abstract

**Background:** Chronic edema (CO) is a progressive, physically disfiguring and currently incurable condition. A multifaceted program has been recommended to manage the swelling. However, there is little evidence investigating patients’ perspectives following the program, particularly for those who have poor adherence or are disengaged.

**Aim:** To investigate the perceived challenges faced by disengaged participants with lower limb CO by identifying their enablers and barriers to participating in a Physiotherapy CO program.

**Method:** An exploratory qualitative approach was used. A purposive sampling strategy was adopted to recruit participants. Those with more than three months swelling and who had low adherence or attendance (disengaged) to the CO program were invited to participate. Semi-structured interviews with six participants from a CO clinic in a tertiary hospital were conducted. Data was thematically analyzed and findings in terms of enablers and barriers were subsequently reflected in the light of a theoretical framework.

**Results:** All six participants were morbidly obese (BMI 47 ± 4 kg/m^2^) with multiple chronic comorbidities. Enablers and barriers detected included physical, psychological and social factors that interplay to present multidimensional challenges that influence the participants’ adjustment to managing their CO. For the disengaged participants in this study, their under-managed lower limb CO was a progression towards being housebound and having a gradually increasing level of disability.

**Conclusion:** Chronic edema, if left unmanaged, can have a profound debilitating impact on the individual permanently. Understanding the challenges faced by patients with lower limb CO has implications for developing a more patient-centered multidisciplinary approach to clinical practice and suggests the need for further research.

---

> *Mentality plays a big part in it. You’ve got to have confidence in the people who are doing things for you. You’ve got to look at it as an optimist, if you’re going to look at it as a pessimist then you are not going to get better. You have to feel like you can keep going (P2)*.

## Introduction

Chronic edema (CO) has been defined as swelling in the tissue that persists beyond three months, and is considered a progressive, physically disfiguring and incurable condition [1, 2]. Primary CO can occur with congenital malformation or secondary to other diseases including obesity, venous disease or cancer treatment [1–3]. Regardless of the etiology, CO has been deemed to be a clinical manifestation of lymphatic system failure [4, 5]. Hence, current opinion recommends CO, rather than “lymphedema”, as a more encompassing term to be used for edema arising from lymphatic origins [6, 7].

Chronic edema is under-recognized and under-managed globally [8]. This has been postulated as due to a lack of published diagnostic and epidemiological data [7, 9, 10]. Hence, there is little published data on prevalence rates of the CO of a non-cancer etiology. A UK epidemiology study (n=823) reported a crude prevalence rate of 1.33/1,000, but this was considered an underestimate due to the methodological parameters [2]. More recently, a subsequent study (n=971) by the same research group reported the crude prevalence rate to be three times higher, at 3.93/1,000 [11]. Prevalence rates have been shown to increase with age, being 10.31/1,000 for those aged between 65 and 74 years and 28.75/1,000 for those aged over 85 years [11]. A gender difference was also demonstrated, with a prevalence rate of 5.37/1,000 in females and 2.48/1,000 in males [11].

Chronic edema is a major clinical problem that impacts on an individual’s physical and psychosocial wellbeing [9, 11, 12]. Unmanaged CO predisposes the individuals to leg ulcers/wounds, recurrent cellulitis and possibly hospitalization [3, 11, 13]. Accordingly, a previous study reported a CO prevalence rate of 28.5 percent amongst hospital inpatients [11]. According to the most recent and largest epidemiology international study on CO, the ‘Lymphedema IMpact and PRevalence INTernational’ (LIMPRINT), that involved collaboration across nine countries and 13,141 participants, there was a higher prevalence rate of 38% (723 patients) amongst acute hospital inpatients [14, 15]. Significant risk factors associated with CO were reported to be advancing age, obesity, immobility and comorbidities including diabetes mellitus [15]. One of the complications of CO involves sustaining wounds commonly in the lower limb impacting on the individual’s mobility [15, 16]. Chronic edema was significantly associated with a history of cellulitis [15], namely that patients with and without wounds suffered at least one event of cellulitis compared to those without CO.

Literature also extensively describes the profound debilitating impact of CO on people’s physical and psychological well-being [17–20]. Compared to their peers without CO, those with the condition present poorer psychological adjustment and lowered health-related quality of life [21]. The impact of CO on health-related quality of life was reported to be worse in those with lower limb edema compared to those with upper limb edema [16] in the aforementioned LIMPRINT study.

Currently, CO is incurable but can be managed. A multifaceted program consisting primarily of compression therapy has been recommended as best practice [5, 22–24]. When recommendations within the program are consistently adhered to, the positive outcomes anticipated are i) a successful reduction in swelling, ii) decreased morbidity, and iii) improved functional outcomes [25]. Despite the benefits, not all patients engage in or adhere to the recommendations in the clinical program for management.

Adherence to therapy and recommendations for a long-term condition are considered essential but described as a complex multidimensional challenge [26–29]. Adherence is beneficial for patients’ health and has economic benefits [29]. Albeit in another area of research, a systematic review of compression therapy for people with leg ulcers reported low adherence rate and recommended further research to investigate patients’ perspectives and factors impacting on adherence [30]. Trust in the health professional was found to be pivotal for leg ulcer treatment [31]. Suggestions to investigate coping strategies or trial cognitive coaching have been proposed as measures to improve adherence amongst individuals with CO [19, 32, 33]. However, there is relatively little evidence on enablers or barriers specifically related to engagement and adherence with a CO program [9, 20, 34] or indeed little application of a theoretical framework in understanding the challenges faced. Using a theoretical framework to evaluate a complex challenge has been recommended [35, 36]. An established theory offers a coherent evidence-based insight into the influences of health behavior and can elucidate mediating factors or strategies towards behavior change in implementation approach later [35, 36].

Therefore, an in-depth investigation into perspectives of individuals with CO who are non-adherent or disengaged to a CO management program, based on their unique lived experience and reflected using a theoretical framework, is required [37]. Subsequently, the understanding of the challenges people with CO face during a CO program and the factors that influence their adherence or non-adherence to the program, will contribute to the development of more relevant and patient-centered CO program, that is better attuned to individual’s needs.

### Aim of the study

The primary aim of the pilot study is to explore the perceived challenges faced by disengaged patients with lower limb CO through the framework of enablers and barriers to participation in a physiotherapy CO program.

## Methodology

### Design

The research used a concurrent/convergent mixed method research approach. This involved the use of qualitative and quantitative approaches concurrently within the study [38]. A mix of these approaches facilitated triangulation of the findings and importantly, provided insights to enhance comprehensiveness in understanding participants’ perspectives [39]. This paper will report the qualitative findings.

### Chronic edema program setting

This project was undertaken at a teaching hospital located in metropolitan Perth in Western Australia. It is a 607-bed public teaching hospital. The Lower Limb Lymphedema Outpatient Clinic (herein, the clinic) is a specialty fee-free service within the hospital’s physiotherapy department operating since 1993. This service currently involves three part-time physiotherapist staff, but no medical doctor. Of the three staff members, two possess postgraduate qualifications and are fully trained therapists in management of CO. The clinic operates as a four-day weekly service with an approximate caseload of 80 patient with consultations totaling in 550 occasions of service annually. Most of the referrals are from medical specialists or allied health personnel. Patients are eligible for treatment if they reside in the hospital catchment area and/or are managed by a doctor within the hospital.

The CO program offered includes assessment and best practice management. [23, 40]. With the development of new technologies over time, the clinic has expanded to offer a broad range of clinical services for CO management. Strategies are primarily based on a) compression therapy (e.g. multi-layered bandaging, compression stockings, compression wraps and stiff self-adhesive bandages e.g. 3M™ Coban™ 2 Compression System); b) education on self-management strategies including manual lymphatic drainage massage and exercise; and c) referral to podiatry and other relevant referral, and/or for further investigations.

### Recruitment

A purposive sampling strategy, specifically, “Homogeneous Sampling” was used to recruit participants [41]. This involved selecting a sub-group of current or previous patients from the clinic, who shared common characteristics and had the potential to provide information-rich data for an in-depth understanding of the research inquiry. The inclusion criteria were:

- Lower limb edema of more than three months duration
- Recurrent presentations/referrals to the clinic
- Disengagement (i.e. low adherence) with the program, including one or more of the following:

- Attendance to ≤ 50% of scheduled clinic appointments, or not attending (disengaged)
- Low uptake or adherence to prescribed therapy recommendations without valid reasons
- Self-reported low compliance or adherence to recommendations

Criteria to exclude individuals from participating were:

- Active malignancy (e.g. undergoing active treatments such as radiotherapy or chemotherapy).
- Insufficient English language competency.

### Data collection and procedure

Potential participants (i.e. those who met the eligibility criteria) were identified by the team’s physiotherapists and contacted to ascertain their interest in being involved in the study. Participants who agreed to take part in the study were offered to choose their preferred venue for the interview. Subsequently, interested participants were interviewed and provided socio-demographic details including information related to their condition. Data on edema type, chronicity of lower limb edema and source of referral were also collected.

### Ethical considerations and approvals

Ethical and governance approval were sought and granted by the hospital’s Human Research Ethics Committee (RGS0000000706). This included permission to access the participants’ relevant medical records and costings. All participants were given a copy of the Participant Information Sheet and subsequently provided written consent.

## Materials

### Interviews

Face-to-face, semi-structured interviews were conducted by the primary researcher (LK). The recorded interview was guided by an interview guide and additional field notes were taken.

The interview guide was derived from previous literature [20, 42–44] and expert advice. Questions and prompts during each interview were tailored to describe and facilitate the participant’s reflections and lived experience, specifically seeking information on their experience with the program in the clinic (e.g. treatment, care received, management strategies outside of the clinical setting and perceived progress of their condition) and personal considerations (perceived assessment of management success, challenges, control over their condition and subsequent impacts and potential changes).

### Theoretical framework COM-B model linked with TDF

The COM-B model is an acronym for Capability, Opportunity and Motivation factors that may affect Behavior. The model posits that behavior is influenced by these three core constructs and any change in health behavior will be dependent upon associated mediating factors [45]. The COM-B model’s constructs are:

- Capability – (Physical): Physical capacity to perform the behavior e.g. skill or strength;
- Capability – (Psychological): Psychological capacity to perform the behavior (e.g. knowledge or comprehension);
- Opportunity – (Physical): Opportunity afforded by the environment involving physical ‘affordance’ and resources;
- Opportunity – (Social): Opportunity afforded by interpersonal influences, social cues and cultural norms that influence the way that we think about things;
- Motivation – (Reflective): Reflective processes involving plans (self-conscious intentions) and evaluations (beliefs about what is good and bad);
- Motivation – (Automatic): Automatic processes involving emotional reactions, desires (wants and needs), impulses, reflex responses.

Another theoretical model Theoretical Domains Framework (TDF) is an evidence-based framework that has several domains encompassing behavior change techniques (Table 1) including affective and emotive constructs [46]. The TDF complements and expands the COM-B model to identify and evaluate techniques of behavior change for application or later implementation. The COM-B and TDF approaches have been applied in a wide range of healthcare research such as smoking cessation behavior, health professionals’ adherence to low back pain guidelines, and falls prevention program [47–49]. In the current study, the COM-B model is mapped against the TDF domains (Table 1) to aid interpretation of findings.

**Table 1.**
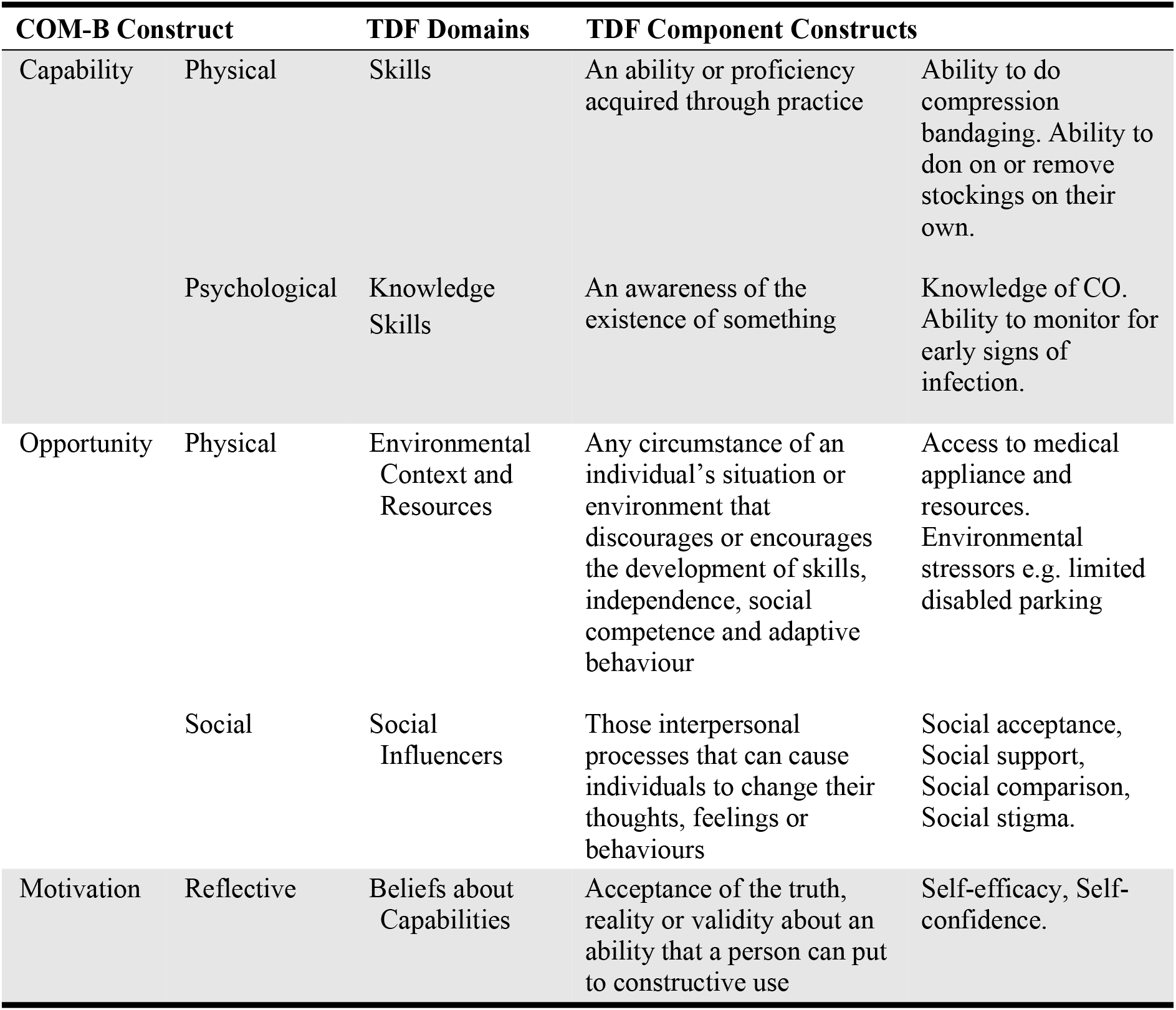
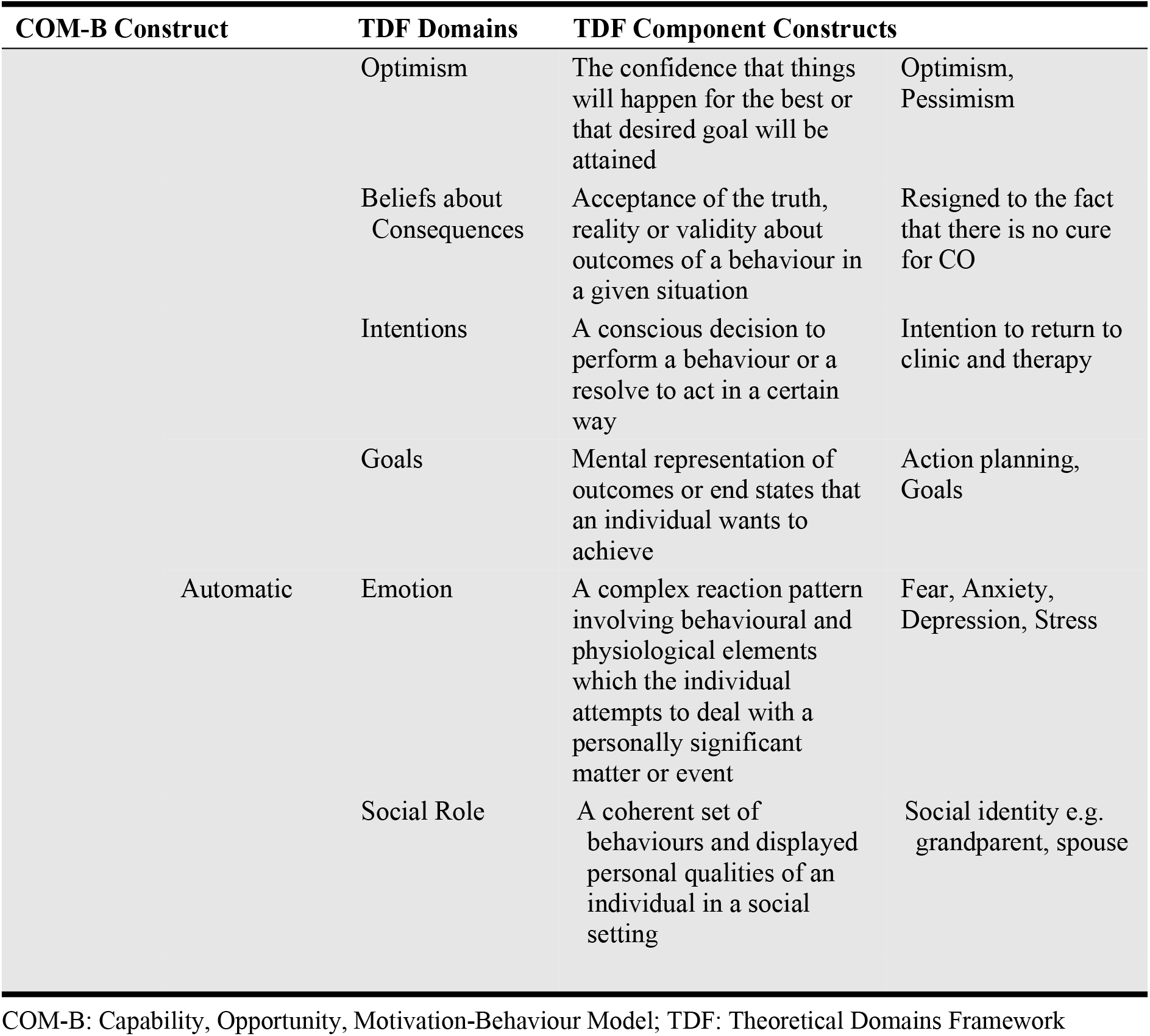
Mapping of the COM-B model to the TDF constructs relevant to the study

### Data analysis

All qualitative data were de-identified and transcribed before importing into NVivo12 [50], a qualitative data management tool to facilitate analysis. Initially the qualitative data were analyzed using thematic analysis [51] through an inductive iterative process whereby text segments with possible common meaning are identified in the transcript, coded and sifted. During this process, attention to the nuances of the dialogue and deeper insights are drawn from the data and emerging themes are proposed [52]. Using investigator triangulation to enhance trustworthiness (credibility) of the findings, a second team member (AB) also examined the data independently, and the emerging themes and areas of agreement /disagreement were discussed with the primary researcher (LK) [51, 53].

Following thematic analysis, the findings were interpreted deductively and reflected in light of the COM-B model and TDF component constructs [45]. Themes and outcomes were determined by consensus between the two researchers prior to sharing with other team members. These were further categorized as enablers or barriers.

## Results

### Characterizing Participants

The sample size (n=6) of this small pilot study was determined by the nature of the patient cohort and the funding grant. Six semi-structured interviews were conducted from September to October 2018. Five interviews were conducted at the participants’ home and one at the clinical setting. Two of the participants’ spouses were present throughout the interview session and were given the opportunity at the end of the session to add further comments.

Participant characteristics are presented in Table 2. Their age ranged between 51 and 72 years, with four (67%) participants living in identified higher socio-economic areas [54]. The participants’ presentation would be deemed as complex cases with more than one decade of CO and with multiple chronic comorbidities since onset of CO. All the participants were morbidly obese, and four (67%) had diabetes. Other characteristics of participants including marital status and details on their medical conditions are provided in Table 2.

**Table 2.**
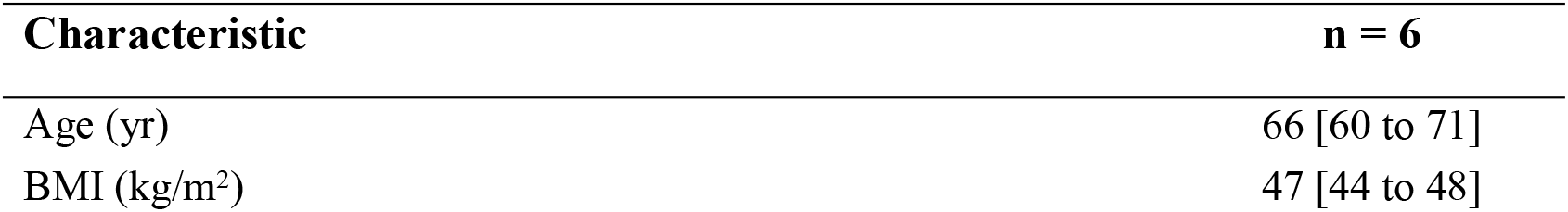
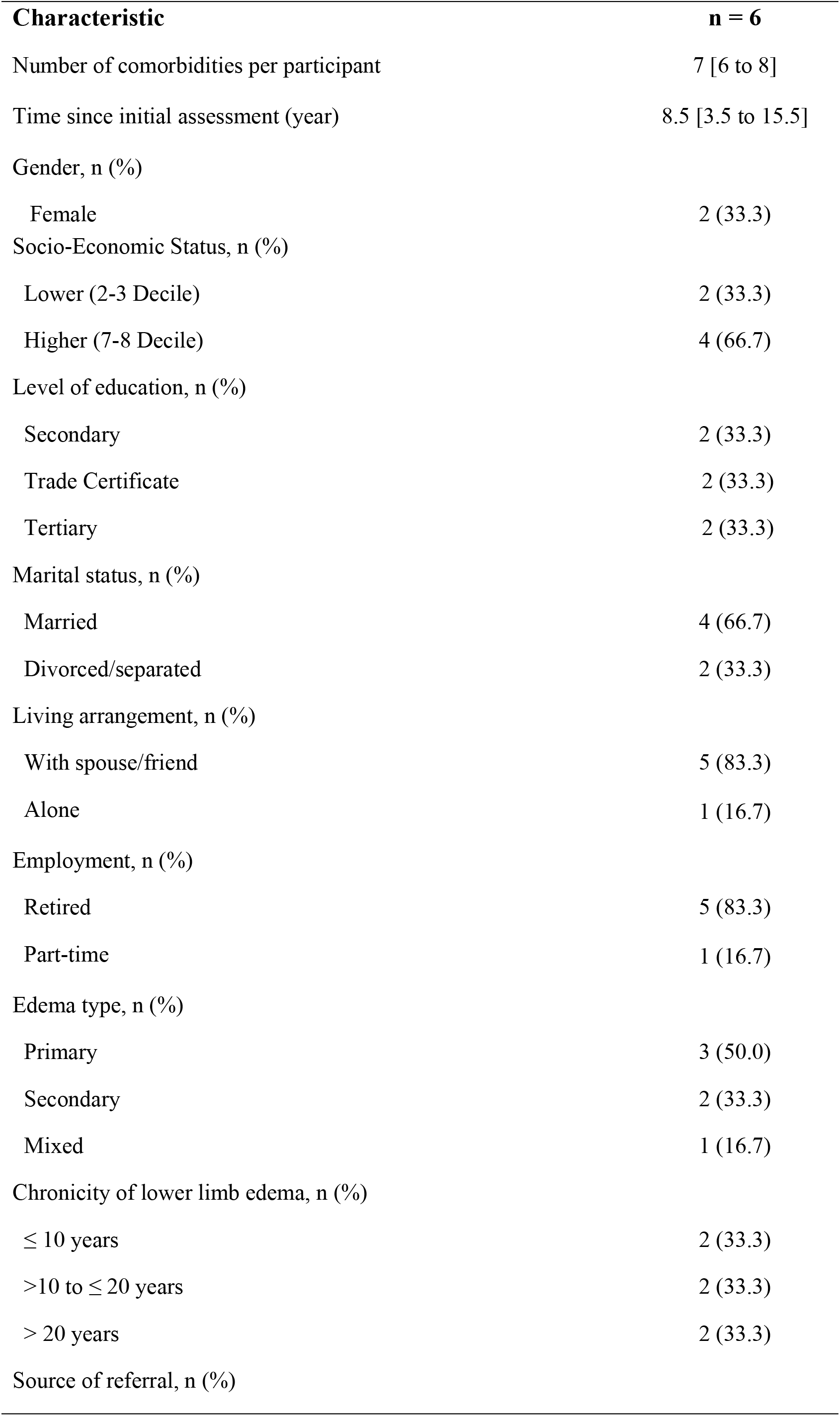
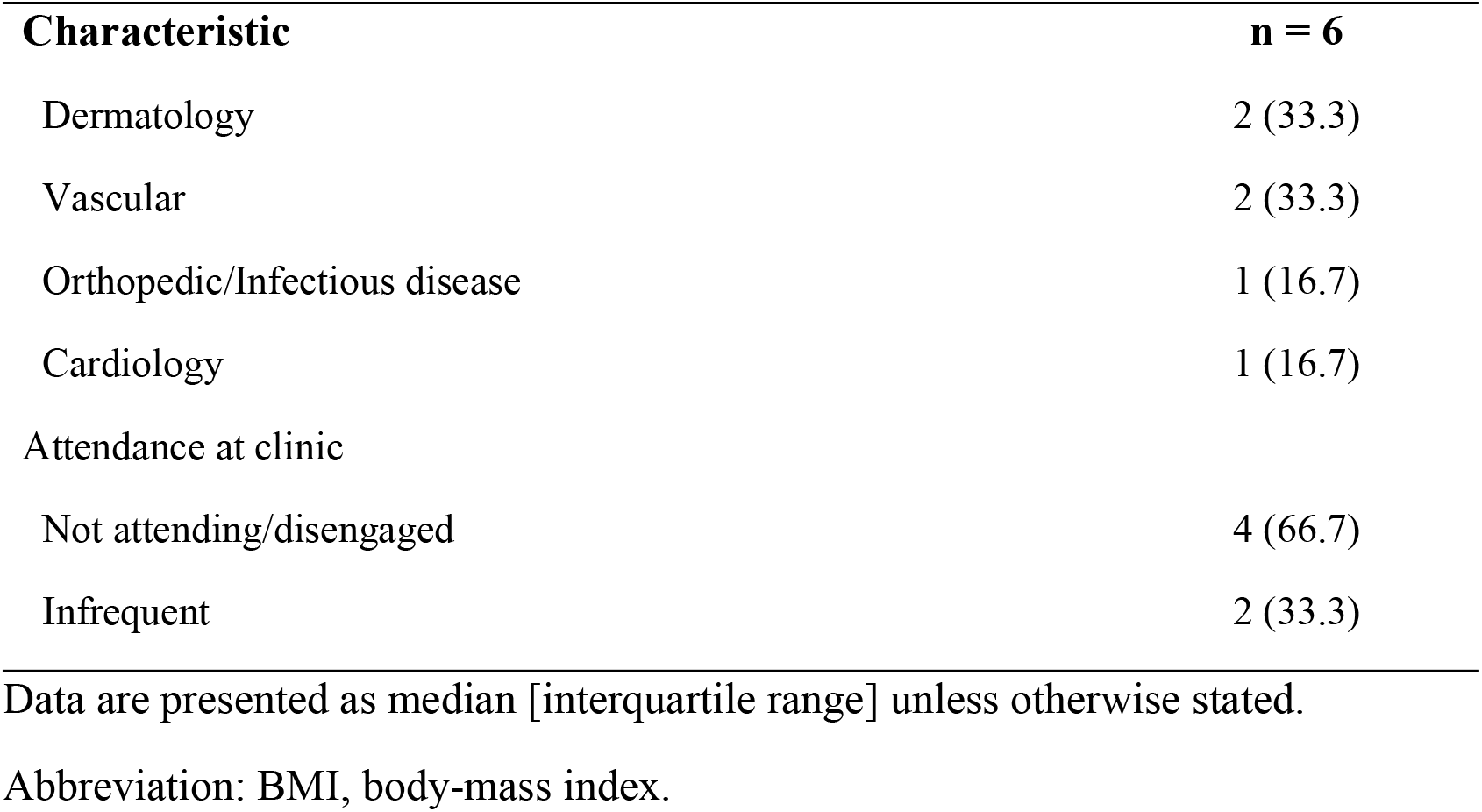
Participants’ characteristics

| Characteristic | n = 6 |
| --- | --- |
| Age (yr) | 66 [60 to 71] |
| BMI (kg/m<sup>2</sup>) | 47 [44 to 48] |

| **Characteristic** | **n = 6** |
| --- | --- |
| Number of comorbidities per participant | 7 [6 to 8] |
| Time since initial assessment (year) | 8.5 [3.5 to 15.5] |
| Gender, n (%) |  |
| Female | 2 (33.3) |
| Socio-Economic Status, n (%) |  |
| Lower (2-3 Decile) | 2 (33.3) |
| Higher (7-8 Decile) | 4 (66.7) |
| Level of education, n (%) |  |
| Secondary | 2 (33.3) |
| Trade Certificate | 2 (33.3) |
| Tertiary | 2 (33.3) |
| Marital status, n (%) |  |
| Married | 4 (66.7) |
| Divorced/separated | 2 (33.3) |
| Living arrangement, n (%) |  |
| With spouse/friend | 5 (83.3) |
| Alone | 1 (16.7) |
| Employment, n (%) |  |
| Retired | 5 (83.3) |
| Part-time | 1 (16.7) |
| Edema type, n (%) |  |
| Primary | 3 (50.0) |
| Secondary | 2 (33.3) |
| Mixed | 1 (16.7) |
| Chronicity of lower limb edema, n (%) |  |
| ≤ 10 years | 2 (33.3) |
| >10 to ≤ 20 years | 2 (33.3) |
| > 20 years | 2 (33.3) |
| Source of referral, n (%) |  |
| Dermatology | 2 (33.3) |
| Vascular | 2 (33.3) |
| Orthopedic/Infectious disease | 1 (16.7) |
| Cardiology | 1 (16.7) |
| Attendance at clinic |  |
| Not attending/disengaged | 4 (66.7) |
| Infrequent | 2 (33.3) |

Data are presented as median [interquartile range] unless otherwise stated.
Abbreviation: BMI, body-mass index.

Thematic analysis identified some further common characteristics not collected as demographics. Over 80 per cent of participants ceased attending the clinic as their condition progressed. Participants frequently described a preference for the management that they initially received in the clinic which included, a period of bandaging prior to being fitted with stockings despite being aware that other treatment options are now available. While the participants adhered to some self-management strategies, one-third of participants with multiple comorbidities chose to manage these conditions at the expense of managing their lymphedema. All participants reported difficulty physically accessing the clinic. There was in general a perceived lack of control over their condition and all participants were subsequently resigned to progressive worsening of lower limb CO, including severe limitations with regard to mobility to the point of being housebound or disabled. All participants were mindful that there is no cure for CO but were on a day-to-day basis dealing with physical, psychological and social impacts. However, despite these, no participants in the study mentioned their weight (obesity) in relation to CO.

### Applied model constructs

In the COM-B model, the first construct is capability both physical and psychological, each of these have enablers (positive factors) and barriers (negative factors) (see Table 1 and Table 3) that were examined in the study.

**Table 3.** Enablers and barriers to adherence to the CO program as identfied and reflected by the COM-B model and the TDF component constructs

| Enablers | Barriers | Examples |
| --- | --- | --- |
| **Capability- Physical**<br>None reported | **Capability- Physical**<br>Decreased physical skills<br>Multiple chronic co-morbidities | Decreased capacity to walk<br>Fear of falling<br>Loss of independence<br>Obesity, diabetes, etc. |
| **Capability- Psychological**<br>Knowledge/procedural knowledge on self-management strategies e.g. bandaging | **Capability- Psychological**<br>Lack of practical knowledge by medical practitioners | Reduced confidence in GP to deal with their infections |
| **Opportunity- Physical**<br>Access to medical appliance, aids and latest options<br>Tailored program to individual's needs and capability | **Opportunity- Physical**<br>Difficulty access to hospital clinic physically<br>Limited appropriate personal resources | Limited disabled parking<br>Limited hospital internal transport service<br>Bus stop is too far from clinic<br>Limited choice of safe fitting footwear and clothes |
| **Opportunity- Social**<br>Positive health provider-patient relationship<br>Continuity of care<br>Partner's social support | **Opportunity- Social**<br>Social stigma<br>Social pressure<br>Social comparison |  |
| **Motivation-Reflective**<br>Belief that therapy and medical appliance helps to reduce CO | **Motivation-Reflective**<br>Pessimism<br>Belief that there is no cure<br>Lack of Motivation, Intention and Goals | Fatalistic attitude<br>Resignation, Resilience<br>Diminished self-efficacy |
| **Motivation-Automatic** | **Motivation-Automatic** |  |
| To feel competent in social role<br>Strive to stay positive | Emotion | Fear, Hurt, Depression,<br>Anxiety |

COM-B: Capability, Opportunity, Motivation-Behaviour Model; TDF: Theoretical Domains Framework

### Capability-Physical

No physical capability enablers were identified, indicating that the long-term impact of CO is debilitation, namely lack of or reduced physical capacity (Table 3). Participants, however, identified numerous barriers including a reduced physical capacity to walk and move due to lower limb CO and a heightened fear of falling. These factors negatively impacting on their mobility, independence, lifestyle choices, self-efficacy and ultimately their quality of life. The concern with severe physical limitation as a barrier was prevalent throughout the study. “The biggest trouble I’m finding with the legs is that they feel heavy and I can’t walk a long way now” (P6). Everyday activities such as grocery shopping were reported to be challenging. This is objectively reflected by a spouse of one of the study participants:

> She can only walk 50m. She gets pain and sits down. We find it easy to go to places with free wheelchairs … it also means her life is pretty much housebound whether she likes it or not. We have to hire a wheelchair to go to the zoo, go to the shops or anything like that (P4’s spouse).

Over time, there was a growing need, and sometimes unease, with the use mobility aids (e.g. wheelchair, or walker). One of the participants (P2) expressed frustration with “I hate going in a wheelchair”. Ambivalence about using mobility aids was commonly discussed:

> I am just too young [50 years old] for a walker (P4).
>
> I don’t feel so steady on my legs anymore and I feel like I am going to fall over. And if I fall over, my knees are so sore I can’t kneel to get up. If I am down, how am I going to get up? So the walker does help get a bit more confidence walking with it (P3).

As identified in the previous comment, a fear of falling was raised throughout the study. Hence, the participants have adapted or altered their daily activities, or even stopped doing what they used to do. As one described about his daily routine as “Because you can’t move, mine goes from sitting to sleeping, then sleeping to sitting” (P5). Others demonstrated how CO can be such a debilitating condition in the long term, leading to a loss of independence and physical capacity. “I can’t lift my legs so my friend comes in everyday and helps me to get into bed” (P2) and “Things like playing tennis. I know that’s impossible now. I just can’t do it” (P6).

Unsurprisingly, given their physical limitations from CO and multiple comorbidities (including obesity), these participants struggled to apply or use the medical appliances and aids (e.g. stockings provided to manage their lower limb CO). P3 described in her earlier years of experience with stockings “very hard to put it on…I persevered for a few months but it is hard”. This perspective has a psychological aspect, which points to another construct (capability-psychological) of the COM-B model.

### Capability-Psychological

Enablers and barriers associated with psychological capability as outlined in Table 3 relate to knowledge and awareness in the study were related to attendance at the CO clinic. These translated into comprehending and applying knowledge of CO, self-management strategies (e.g. bandaging, massage, wound care and exercise), monitoring for infection, and requirements of ongoing care of their legs. In contrast, a barrier expressed by all study participants was the lack of practical knowledge and awareness by medical practitioners on ways to treat their CO and on referral pathways, leaving them in most cases solely reliant on the clinic.

### Opportunity-Physical

This COM-B model construct looks at opportunities that are afforded by environmental context, resources and social influences (Table 1 and Table 3). There were more barriers than enablers identified.

Physical enablers discussed by all the study participants included that the CO treatment program was tailored to their needs and preferences and that they had access to most medical appliance options. Barriers identified in the interviews included physically accessing the hospital clinic. Those who drove to the hospital mentioned that it was a challenge to access hospital disabled parking due to limited availability. Those who used public transport reported that the distance between the bus stop and the clinic was too long and walking that far was not easy. Another identified barrier discussed by few participants was their difficulty in finding appropriate clothing and safe footwear when they ventured outside of their house.

### Opportunity-Social

Social influencers including a positive health provider-patient relationship, continuity of care, and a supportive spouse/partner were important enablers suggested. All the participants acknowledged the clinic and health professionals’ expertise in identifying and addressing their CO, providing a flexible and tailored solutions. This was particularly relevant when study participants were actively engaged and adhering to recommendations at the clinic. In addition, having the same health professional in attendance at the clinic over time was reported to be “psychologically relieving” (P5).

Social stigma was identified as a barrier to adherence to therapy. Fear of being ridiculed when wearing medical appliance in public such as “He’s just wearing bandages so that he can get a seat [comments by other passengers in the bus]” (P6) was recounted by one study participant. Associated with these comments was the frustration about “having to explain it [his condition] over and over again” (P6) to acquaintances. This effort could reflect an act of dealing with social pressure and maintaining social acceptance (Table 1 and Table 3).

### Motivation-Reflective

The third construct of the COM-B model is motivation, which includes reflective and automatic processes as well as associated enablers and barriers. Reflective aspects relate to beliefs and intentions while automatic responses entail unconscious desires and impulses.

Whilst participants expressed beliefs that therapy helped with reducing swelling in their legs, their pessimism and resignation towards recurrent setbacks were barriers to their well-meant intentions. Therapy helped to reduce the size of their legs and aided them to continue to function (i.e. attending work, socializing and so forth). This was largely related to persistently wearing stockings in the early years of the condition.

Over time, with their reduced physical capability, their sense of self-efficacy was gradually eroded and evoked a sense of resignation with an increasing reliance on their spouse and family. P6 vividly described this:

> My self-confidence is diminished due to my physical limitations when previously I walked a lot without tiring... I don’t feel safe driving because I don’t feel I can lift my legs to get to the pedals [brake] quickly enough if someone came in front of me. She [spouse] always has to do the driving and even when she is not feeling up to it [unwell]. I can’t help her with that because I don’t feel comfortable.

All participants expressed a lack of control and resilience over their condition because they did not see a cure. Consequently, highly reported was a degree of resignation to just accept the condition and live with it.

> I’m controlling it to the degree, where I don’t have to stay in bed all day. Like I control it to the point where I live a life but I can’t control it to a point where I’m going to cure it or I’m going to get better. I don’t know whether it’s possible to live that…There is no cure so there is nothing you can do (P1)

“I’d say I tolerate it [CO] as I don’t have any choice. The legs haven’t really got any worse but it is hard to say I have got control of it” (P6). Consequently, motivation towards therapy, including attending the clinic and accepting new or alternative treatment options declined.

### Motivation-Automatic

There was a gamut of emotive enablers and barriers displayed that were less conscious. On the one hand, participants expressed a compulsion to feel competent in fulfilling their social role with their family (e.g. as a grandparent, or spouse; striving to stay positive) and a yearning for independence was expressed.

> When you are doing things for yourself, it’s a big plus (P2).
>
> I’d rather do it myself [drive to the hospital]. I would still like to be independent (P3).
>
> My biggest thing is I like to be independent and keeping everything under control makes me happiest because I am not looking for great outcomes or anything except maintenance. Maintaining what I’ve got now (P5).

On the other hand, having CO over a long period has had both a reported physical and an emotional effect on study participants. While the physical impacts have been explored throughout, for other participants its impact was an inhibitor, namely that CO “changed my whole state of mind” (P2). Emotions expressed and detected from the interviews included emotional pain, depression, fear and anxiety; they were seen as having long term impact. “Seeing something visually can affect you emotionally … Hurt, I look and think I used to [be] called elephant legs when I was in school” (P4).

Interestingly, utterances by the male participants were inclined to talk more often about “the legs” and rarely as “my legs”. In addition, weight was associated with the legs and not associated with their body weight despite participants having higher than average Body Mass Index (BMI) (Table 2). Hence, these appeared to be a coping strategy with a disembodiment of their legs and body weight in their comments. In relation, diet was not uttered during the interviews as a strategy to manage their CO. Only one of the six participants had current obvious intention to manage their CO and mentioned goal-setting and CO action plan.

## Discussion

Low adhering or disengaged patients have broadly been deemed as non-compliant [29], however, the findings of this pilot study have shown that can be an oversimplification. By exploring participants’ perspectives, this study has revealed multifactorial issues (enablers and barriers) that complicate adherence and attendance at the CO clinic. Further, the progressive nature of lower limb CO, compounded by patient disengagement from therapy inflicted a profound, long-term, debilitating effect on this small cohort.

Evident in the pilot study was an overwhelming negative impact of lower limb CO on quality of life, independence, psychological distress, expectations, as well as resignation and inevitable permanent disability of participants in their later years. Despite adherence to treatment and self-management strategies at an early stage following diagnosis, managing lower limb CO long term has been challenging. Findings from this study were consistent with other research [18, 20, 55]. In an Irish study, data was collected using a questionnaire, the cohort was younger (average 54 years old) with the majority living in the countryside [18]. That study supports the findings of this pilot in establishing that CO impacts on social functioning and leads to social isolation. Findings in two other interview-based studies conducted in UK [20] and Israel [55] were similar to this pilot study. Tidhar and Armer (55) found that while participants related success with no exacerbation in the CO, their expectations of “getting better” faded over time. Watts (2016) also pointed to the management of CO as challenging due to physical and functional constraints being entangled with psychosocial impacts, including social stigma. However, the absence of reporting on age of the participants, BMI profiles and the length of time with the condition limited the ability to compare characteristics with the sample cohort. While these studies described various consequences of CO, none looked beyond narrative nor towards an analytical identification of the challenges using a theoretical framework.

The current study has applied the theoretical framework COM-B model linked with TDF (Table 1) in understanding capability, opportunity and motivation factors as the lens for interpreting patient disengagement in therapy via enablers and barriers. The study found that the enablers and barriers encompass physical, psychological and social factors impacting on the participants’ adherence to therapy. These factors entwined as complex multidimensional challenges that impacted participants adjusting to living with a chronic condition [29, 56, 57]. Importantly, some of the factors were internal (personal realm) and others as external factors beyond their control. On the personal level, having a supportive partner was found to positively facilitate communal coping [57] towards adjusting to CO. Externally, the major barrier for these participants was a lack of capacity to access the hospital physically to seek therapy. This is consistent with the LIMPRINT study where nearly one-third of the participants (35.5%) reported distance as preventing access to treatment [16]. This is a cohort of individuals with multiple chronic conditions, including unhealthy weight. Obesity is not necessarily unique to this cohort, as it has estimated that 63.4 per cent of Australian adults are considered overweight [58]. There is growing evidence that being obese is a significant risk factor for CO [15, 59, 60] however it was infrequently highlighted by the study cohort.

Efforts to facilitate behavior change in people with lower limb CO would benefit from being guided by a health behavior change theoretical framework(s) [61]. This would heighten the appreciation and learning from current practice in people with other chronic conditions, such as those with cardiac disease, or obesity [62]. Participants were keen to participate and be heard, their anecdotes indicated limited awareness of appropriate management of CO amongst the medical practitioners; a finding that is consistent with studies conducted in Canada, USA, UK and the global LIMPRINT study [9, 63, 64].

Tinetti and colleagues Tinetti, Green (65) urged patient-centered care for the cohort of people with multiple comorbidities by advocating that health providers treat the individual, and not the disease, through tailoring care to the individual’s priorities, rather than strictly following practice guidelines. The current study highlights the need for ongoing patient-centered care. Understanding the contextual factors, as illustrated by the enablers and barriers, influencing the individual’s adjustment to and management of chronic illness [56], can be an important first step towards successful management of lower limb CO.

The current study is not without limitations. The sample size is small; however, people with lower limb CO are a “hard-to-reach” population [20, 66]. We believe that the sample for our exploratory qualitative study reflects many of the findings from other empirical studies as identified. The views of a marginalized cohort of mostly older people with CO who are housebound and disengaged have been reported [67]. In that sense, the findings reflect the experience of a purposive sample of participants from particular clinic in a metropolitan area of Western Australia; and therefore, caution should be taken when trying to generalize our findings.

## Conclusion

This study has identified enablers and barriers faced by low adherent or disengaged participants with lower limb CO to participating in a hospital-based CO program. This study has also provided insights into the multidimensional challenges faced by these participants. Findings revealed that CO, if under-managed, can have a profound debilitating impact on the individual and may be a progression towards inevitable permanent disability. The findings have implications for a more patient-centered clinical practice, indications for a multidisciplinary approach and further research.

## Author contributions

Conceptualization: LK

Design: LK, AB, WJ, VC

Ethical process: LK, VC

Recruitment: LK, WJ

Interviews, data collection, data transcription, qualitative data analysis: LK, AB

Quantitative data analysis: LK, VC

Preparation and final approval of manuscript: LK, AB, WJ, VC

## Funding

This study (LK) was supported by the Sir Charles Gairdner Hospital Research Advisory Committee’s Small Grant 2017. The funding organization had no role in the study design, data collection and analysis nor in preparation of the manuscript. VC is supported by Cancer Council of Western Australia Postdoctoral Research Fellowship.

## Competing Interests

The authors have declared no competing interests.

## Acknowledgements

We would like to thank the following people for their help:

- Sir Charles Gairdner Hospital Physiotherapy Department staff, in particular Alison Williams (retired) and Patricia Lobo for their help with recruitment of participants.
- Royal Perth Hospital Library staff especially Rina Rukmini for her help and support with literature search.
- Patients and therapists who contributed to the questionnaire and interview guide.
- LK Lymphoedema Centre for contributing some of the fund for publishing as open access.

## Data Availability Statement

Due to ethical considerations protecting participant and hospital confidentiality, the interview data are not publicly available.

## Reference

1. Keeley V. Advances in understanding and management of lymphoedema (cancer, primary). Current Opinion in Supportive and Palliative Care. 2017;11(4):355–60. Epub 2017/10/07. doi: 10.1097/SPC.0000000000000311.

2. Moffatt CJ, Franks PJ, Doherty DC, Williams AF, Badger C, Jeffs E, et al. Lymphoedema: An underestimated health problem. QJM. 2003;96(10):731–8. doi: 10.1093/qjmed/hcg126. PubMed PMID: 14500859.

3. Szuba A, Rockson SG. Lymphedema: classification, diagnosis and therapy. Vascular Medicine. 1998;3(2):145–56. doi: 10.1177/1358836X9800300209.

4. Mortimer PS, Levick JR. Chronic peripheral oedema: The critical role of the lymphatic system. Clin Med (Northfield Il). 2004;4(5):448–53.

5. Stout N, Partsch H, Szolnoky G, Forner-Cordero I, Mosti G, Mortimer P, et al. Chronic edema of the lower extremities: International consensus recommendations for compression therapy clinical research trials. Int Angiol. 2012;31(4):316–29.

6. Moffatt CJ, Doherty DC, Franks PJ, Mortimer PS. Community-based treatment for chronic edema: An effective service model. Lymphat Res Biol. 2018;16(1):92–9. doi: 10.1089/lrb.2017.0021.

7. Moffatt C, Keeley V, Quere I. The concept of chronic edema-A neglected public health issue and an international response: The LIMPRINT Study. Lymphat Res Biol. 2019;17(2):121–6. doi: 10.1089/lrb.2018.0085.

8. Rockson SG. LIMPRINT: Elucidating the global problem of lymphedema. Lymphat Res Biol. 2019;17(2):119–20. doi: 10.1089/lrb.2019.29062.sr.

9. Keast DH, Despatis M, Allen JO, Brassard A. Chronic oedema/lymphoedema: under-recognised and under-treated. International Wound Journal. 2015;12(3):328–33. doi: 10.1111/iwj.12224.

10. Nairn S, Dring E, Aubeeluck A, Quere I, Moffatt C. LIMPRINT: A sociological perspective on “Chronic Edema”. Lymphat Res Biol. 2019;17(2):168–72. doi: 10.1089/lrb.2018.0082.

11. Moffatt CJ, Keeley V, Franks PJ, Rich A, Pinnington LL. Chronic oedema: A prevalent health care problem for UK health services. International Wound Journal. 2017;14(5):772–81. doi: 10.1111/iwj.12694.

12. Morgan PA, Murray S, Moffatt CJ, Honnor A. The challenges of managing complex lymphoedema/chronic oedema in the UK and Canada. International Wound Journal. 2012;9(1):54–69. doi: 10.1111/j.1742-481X.2011.00845.x.

13. Damstra RJ, van Steensel MA, Boomsma JH, Nelemans P, Veraart JC. Erysipelas as a sign of subclinical primary lymphoedema: A prospective quantitative scintigraphic study of 40 patients with unilateral erysipelas of the leg. Br J Dermatol. 2008;158(6):1210–5. doi: 10.1111/j.1365-2133.2008.08503.x.

14. Moffatt C, Franks P, Keeley V, Murray S, Mercier G, Quere I. The development and validation of the LIMPRINT methodology. Lymphat Res Biol. 2019;17(2):127–34. doi: 10.1089/lrb.2018.0081.

15. Quere I, Palmier S, Noerregaard S, Pastor J, Sykorova M, Dring E, et al. LIMPRINT: Estimation of the prevalence of lymphoedema/chronic oedema in acute hospital in In-Patients. Lymphat Res Biol. 2019;17(2):135–40. doi: 10.1089/lrb.2019.0024.

16. Mercier G, Pastor J, Moffatt C, Franks P, Quere I. LIMPRINT: Health-related quality of life in adult patients with chronic edema. Lymphat Res Biol. 2019;17(2):163–7. doi: 10.1089/lrb.2018.0084.

17. Fu MR, Ridner SH, Hu SH, Stewart BR, Cormier JN, Armer JM. Psychosocial impact of lymphedema: A systematic review of literature from 2004 to 2011. Psychooncology. 2013;22(7):1466–84. doi: 10.1002/pon.3201

18. Greene A, Meskell P. The impact of lower limb chronic oedema on patients’ quality of life. International Wound Journal. 2017;14(3):561–8. doi: 10.1111/iwj.12648.

19. Ridner SH, Shih YCT, Doersam JK, Rhoten BA, Schultze BS, Dietrich MS. A pilot randomized trial evaluating lymphedema self-measurement with bioelectrical impedance, self-care adherence, and health outcomes. Lymphat Res Biol. 2014;12(4):258–66. doi: 10.1089/lrb.2014.0017.

20. Watts TE, Davies RE. A qualitative national focus group study of the experience of living with lymphoedema and accessing local multiprofessional lymphoedema clinics. J Adv Nurs. 2016;72(12):3147–59. doi: 10.1111/jan.13071.

21. Moffatt CJ, Aubeeluck A, Franks PJ, Doherty DC, Mortimer P, Quere I. Psychological factors in chronic edema: A case-control study. Lymphat Res Biol. 2017;15(3):252–61.

22. International Society of Lymphology. The diagnosis and treatment of peripheral lymphedema: 2016 consensus document of the International Society of Lymphology. Lymphology. 2013;49(4):170–84.

23. Lymphoedema Framework. Best practice for the management of lymphoedema. International Consensus. 53 Hargrave Road, London N19 5SH, UK: Medical Education Partnership (MEP) Ltd; 2006. Available from: http://www.woundsinternational.com/pdf/content_175.pdfBest

24. Tzani I, Tsichlaki M, Zerva E, Papathanasiou G, Dimakakos E. Physiotherapeutic rehabilitation of lymphedema: State-of-the-art. Lymphology. 2018;51(1):1–12.

25. Stout NL, Brantus P, Moffatt C. Lymphoedema management: An international intersect between developed and developing countries. Similarities, differences and challenges. Global Public Health. 2012;7(2):107–23. doi: 10.1080/17441692.2010.549140.

26. Bissonnette JM. Adherence: A concept analysis. J Adv Nurs. 2008;63(6):634–43. doi: 10.1111/j.1365-2648.2008.04745.x.

27. Nelson EA, Harrison MB, Canadian Bandage Trial T. Different context, different results: Venous ulcer healing and the use of two high-compression technologies. J Clin Nurs. 2014;23(5-6):768–73. doi: 10.1111/jocn.12105.

28. Van Hecke A, Grypdonck M, Defloor T. A review of why patients with leg ulcers do not adhere to treatment. J Clin Nurs. 2009;18(3):337–49. doi: 10.1111/j.1365-2702.2008.02575.x.

29. World Health Organisation. Adherence to long-term therapies: Evidence for action. Geneva, Switzerland: 2003.

30. Nelson EA, Bell-Syer SE. Compression for preventing recurrence of venous ulcers. Cochrane Database Systematic Review. 2014;(9):CD002303. doi: 10.1002/14651858.CD002303.pub3.

31. Van Hecke A, Verhaeghe S, Grypdonck M, Beele H, Defloor T. Processes underlying adherence to leg ulcer treatment: A qualitative field study. Int J Nurs Stud. 2011;48(2):145–55. doi: 10.1016/j.ijnurstu.2010.07.001.

32. Ridner SH, Fu MR, Wanchai A, Stewart BR, Armer JM, Cormier JN. Self-management of lymphedema: A systematic review of the literature from 2004 to 2011. Nurs Res. 2012;61(4):291–9. doi: 10.1097/NNR.0b013e31824f82b2.

33. Williams AF, Moffatt CJ, Franks PJ. A phenomenological study of the lived experiences of people with lymphoedema. Int J Palliat Nurs. 2004;10(6):279–86. doi: 10.12968/ijpn.2004.10.6.13270.

34. Lasinski BB, Thrift KM, Squire D, Austin MK, Smith KM, Wanchai A, et al. A systematic review of the evidence for Complete Decongestive Therapy in the treatment of lymphedema from 2004 to 2011. Physical Medicine and Rehabilitation. 2012;4(8):580–601. doi: 10.1016/j.pmrj.2012.05.003.

35. Improved Clinical Effectiveness through Behavioural Research Group (ICEBeRG). Designing theoretically-informed implementation interventions. Implemention Science. 2006;1:4. doi: 10.1186/1748-5908-1-4.

36. Rimer B, Glanz K. Theory at a glance. A guide for health promotion practice. 2005. Available from: https://pubs.cancer.gov/ncipl/detail.aspx?prodid=T052.

37. Barlow S, Dixey R, Todd J, Taylor V, Carney S, Newell R. ‘Abandoned by Medicine’? A qualitative study of women’s experiences with lymphoedema secondary to cancer, and the implications for care. Prim Health Care Res Dev. 2014;15(4):452–63. doi: 10.1017/S1463423613000406.

38. Morse JM. Simultaneous and sequential qualitative mixed method designs. Qualitative Inquiry. 2010;16(6):483–91. doi: 10.1177/1077800410364741.

39. Creswell JW, Plano Clark VL. Designing and conducting mixed methods research. 2nd ed. Los Angeles, Calif.: SAGE Publications; 2011.

40. Moffatt C, Partsch H, Schuren J, Quéré I, Sneddon M, Flour M, et al. Compression Therapy: A position document on compression bandaging. 2012. Available from: http://www.soffed.co.uk/lymphorg/wp-content/uploads/2016/03/Compression-bandaging-final.pdf.

41. Patton MQ. Designing qualitative studies. Qualitative research & evaluation methods: Integrating theory and practice. 4th ed. Thousand Oaks, California: SAGE Publications Inc.; 2015. p. 243–326.

42. Dial M, Holmes J, McGownd R, Wendler MC. “I do the best I can:” Personal care preferences of patients of size. Appl Nurs Res. 2018;39:259–64.

43. Holdsworth E, Thorogood N, Sorhaindo A, Nanchahal K. A qualitative study of participant engagement with a weight loss intervention. Health Promotion Practice. 2017;18(2):245–52. doi: 10.1177/1524839916659847.

44. Tidhar D, Katz-Leurer M. Aqua lymphatic therapy in women who suffer from breast cancer treatment-related lymphedema: A randomized controlled study. Support Care Cancer. 2010;18(3):383–92. doi: 10.1007/s00520-009-0669-4.

45. Michie S, van Stralen MM, West R. The behaviour change wheel: A new method for characterising and designing behaviour change interventions. Implementation Science. 2011;6(1):42. doi: 10.1186/1748-5908-6-42

46. Cane J, O’Connor D, Michie S. Validation of the theoretical domains framework for use in behaviour change and implementation research. Implementation Science. 2012;7(1):1–17. doi: 10.1186/1748-5908-7-37.

47. Beenstock J, Sniehotta FF, White M, Bell R, Milne EM, Araujo-Soares V. What helps and hinders midwives in engaging with pregnant women about stopping smoking? A cross-sectional survey of perceived implementation difficulties among midwives in the North East of England. Implementation Science. 2012;7:36. doi: 10.1186/1748-5908-7-36.

48. French SD, Green SE, O’Connor DA, McKenzie JE, Francis JJ, Michie S, et al. Developing theory-informed behaviour change interventions to implement evidence into practice: A systematic approach using the Theoretical Domains Framework. Implementation Science. 2012;7. doi: 10.1186/1748-5908-7-38.

49. Khong LAM, Berlach RG, Hill KD, Hill A-M. Design and development of a theory-informed peer-led falls prevention education programme to translate evidence into practice: A systematic approach. International Journal of Health Promotion and Education. 2018:1–16. doi: 10.1080/14635240.2018.1479650.

50. QSR International Pty Ltd. NVivo Qualitative data analysis software. Version 10. 2012.

51. Miles MB, Huberman AM, Saldana J. Qualitative data analysis: A methods sourcebook. 3rd ed. Thousand Oaks, CA: SAGE Publications, Inc; 2014.

52. Bazeley P. Analysing qualitative data: More than ‘identifying themes’. Malaysian Journal of Qualitative Research. 2009;2(2):6–22.

53. Lincoln YS, Guba EG. Establishing trustworthiness. In: Guba EG, editor. Naturalistic inquiry. Beverly Hills, CA: Sage Publications; 1985. p. 289–331.

54. Australian Bureau of Statistics. Socio-economic indexes for areas-postal areas. Belconnen, ACT: Australian Bureau of Statistics, 2013 ABS catalogue 2033.0.55.001.

55. Tidhar D, Armer J. The meaning of success in lymphoedema management. J Lymphoedema. 2018;13(1):37–42.

56. Helgeson VS, Zajdel M. Adjusting to chronic health conditions. Annu Rev Psychol. 2017;68:545–71. doi: 10.1146/annurev-psych-010416-044014.

57. Helgeson VS, Jakubiak B, Van Vleet M, Zajdel M. Communal coping and adjustment to chronic illness: Theory update and evidence. Pers Soc Psychol Rev. 2018;22(2):170–95. doi: 10.1177/1088868317735767.

58. Australian Bureau of Statistics. Australian Health Survey: First Results, 2011–12 2015 [01 April 2019]. Available from: http://www.abs.gov.au/ausstats/abs@.nsf/Lookup/4364.0.55.001Chapter1002011-12.

59. Mehrara BJ, Greene AK. Lymphedema and obesity: is there a link? Plast Reconstr Surg. 2014;134(1):154e–60e. doi: 10.1097/PRS.0000000000000268.

60. Savetsky IL, Torrisi JS, Cuzzone DA, Ghanta S, Albano NJ, Gardenier JC, et al. Obesity increases inflammation and impairs lymphatic function in a mouse model of lymphedema. American Journal of Physiology Heart and Circulatory Physiology. 2014;307(2):H165–72. doi: 10.1152/ajpheart.00244.2014.

61. Michie S, Atkins L, West R. The behaviour change wheel: A guide to designing interventions. UK: Silverback Publishing; 2014.

62. Mattke S, Mengistu T, Klautzer L, Sloss EM, Brook RH. Improving care for chronic conditions: Current practices and future trends in health plan programs. Rand Health Quarterly. 2015;5(2):3.

63. Moffatt CJ, Gaskin R, Sykorova M, Dring E, Aubeeluck A, Franks PJ, et al. Prevalence and risk factors for chronic edema in U.K. community nursing services. Lymphat Res Biol. 2019;17(2):147–54. doi: 10.1089/lrb.2018.0086.

64. Runowicz CD. Lymphedema: Patient and provider education. Cancer. 1998;83(S12B):2874–6. doi: 10.1002/(SICI)1097-0142(19981215)83:12B+<2874::AID-CNCR42>3.0.CO;2-4.

65. Tinetti ME, Green AR, Ouellet J, Rich MW, Boyd C. Caring for patients with multiple chronic conditions. Ann Intern Med. 2019;170:199–200. doi: 10.7326/M18-3269.

66. Faugier J, Sargeant M. Sampling hard to reach populations. J Adv Nurs. 1997;26(4):790–7. doi: 10.1046/j.1365-2648.1997.00371.x.

67. Watts G. Why the exclusion of older people from clinical research must stop. Br Med J. 2012;344:e3445. doi: 10.1136/bmj.e3445.

